# Cholesterol recognition motifs in the transmembrane domain of the tyrosine kinase receptor family: the case for TRKB

**DOI:** 10.1101/734012

**Authors:** Cecilia Cannarozzo, Senem Merve Fred, Mykhailo Girych, Caroline Biojone, Giray Enkavi, Tomasz Róg, Ilpo Vattulainen, Plinio C Casarotto, Eero Castrén

**Author notes:** corresponding author: Plinio Cabrera Casarotto, PhD., Neuroscience Center., Haartmaninkatu 8, B233b. PO Box 63., University of Helsinki, Finland.

## Abstract

Cholesterol is an essential constituent of cell membranes. Recently, the discovery of cholesterol recognition amino acid consensus (CRAC) on proteins indicated a putative direct, non-covalent interaction between cholesterol and proteins. In the present study, we evaluated the presence of a CRAC motif and its inverted version (CARC) in the transmembrane region (TMR) of the tyrosine kinase receptor family (RTK) in several species using *in silico* methods. CRAC motifs were found across all species analyzed, while CARC was found only in vertebrates. The tropomyosin-related kinase B (TRKB), a member of the RTK family, is a core participant in the neuronal plasticity process and exhibits a CARC motif in its TMR. Upon recognition of the conserved CARC motif in the TRKB, we performed molecular dynamics simulations with the mouse TRKB TMR sequence, which indicated that cholesterol interaction with the TRKB CARC motif occurs mainly at the central Y433 residue. Binding assay suggested a bell-shaped effect of cholesterol on BDNF interaction with TRKB receptors. Therefore, CARC/CRAC motifs may have a role in the function of the RTK family TMR.

## Introduction

The human brain contains 23% of the body’s total cholesterol. Most of this cholesterol is found in the myelin sheath of oligodendrocytes (Dietschy & Turley, 2004; Martin *et al.*, 2014). As the blood-brain barrier prevents lipoprotein or cholesterol transport to the brain, local *de novo* synthesis takes place. In the mouse brain, cholesterol synthesis peaks during the second postnatal week and then decreases significantly independent of sex or blood cholesterol concentration (Quan *et al.*, 2003; Pfrieger & Ungerer, 2011). During early development, neurons produce cholesterol autonomously (de Chaves *et al.*, 1997; Nieweg *et al.*, 2009; Pfrieger & Ungerer, 2011). In later stages, cholesterol is synthesized by glial cells. However, it is unknown if this synthesis is constant or under regulated production (Saito *et al.*, 2009; Pfrieger & Ungerer, 2011).

Cholesterol can be localized on both leaflets of the plasma membrane (Fantini *et al.*, 2016) and induces changes in physical properties of the membrane, such as fluidity (Maguire & Druse, 1989) and curvature (Lee, 2004). Cholesterol can also interact with transmembrane domains to regulate protein function (Fantini & Barrantes, 2013; Elkins *et al.*, 2018). Cholesterol is a core constituent of microdomains known as lipid rafts, which serve as signaling platforms for several pathways (Lang *et al.*, 2001; Pereira & Chao, 2007; Zonta & Minichiello, 2013). In the nervous system, cholesterol interaction with membrane proteins influences several crucial events, such as exocytosis of synaptic vesicles (Linetti *et al.*, 2010), synaptic activity, connectivity, plasticity, signal transduction, transmission, and cell survival (Michikawa & Yanagisawa, 1999; Goritz *et al.*, 2005; Liu *et al.*, 2010).

The tropomyosin-related kinase receptor (TRK) subfamily is one of the most prominent subfamilies of tyrosine kinase receptors (RTK) and plays a crucial role in neuronal plasticity (Dekkers *et al.*, 2013). The TRK receptors consist of three members (TRKA, TRKB, and TRKC, or NTRK1, NTRK2 and NTRK3, respectively) that are phosphorylated on several tyrosine residues on the intracellular portion upon activation by their high-affinity ligands (NGF, BDNF, and NT-3, respectively) (Huang & Reichardt, 2001). TRKA and TRKC are located in lipid rafts, while the presence of TRKB in rafts occurs transiently upon BDNF stimulation (Suzuki *et al.*, 2004, 2007). Functionally, in the absence of ligand, TRKA and TRKC (but not TRKB) induce cell death mediated by interaction with p75NTR, the low-affinity receptor of several neurotrophins (Nikoletopoulou *et al.*, 2010; Dekkers *et al.*, 2013).

*In silico* models suggest that two cholesterol molecules can interact in a tail-to-tail fashion as a transbilayer dimer (Harris *et al.*, 1995; Rukmini *et al.*, 2001) or back-to-back through their flat alpha faces, leaving the beta sides accessible for interactions with proteins (Hanson *et al.*, 2008). On these target proteins, the following two consensus motifs with predictive value have been defined (Di Scala *et al.*, 2017): the Cholesterol Recognition Amino acid Consensus sequence (CRAC) and its “inverted” version (CARC) (Li & Papadopoulos, 1998; Baier *et al.*, 2011). The CRAC linear sequence, from N- to C-terminus, consists of an apolar residue (leucine [L] or valine [V]), one to five amino acids of any kind, an aromatic amino acid (tyrosine [Y] or phenylalanine [F]), one to five amino acids of any kind, and a basic residue (arginine [R] or lysine [K]) (Fantini & Barrantes, 2013). CARC consists of the same pattern in the opposite direction, with tryptophan (W) as alternative aromatic residue. CARC has a higher affinity for cholesterol than CRAC (Fantini & Barrantes, 2013; Di Scala *et al.*, 2017). Several proteins have been identified to contain CRAC/CARC motifs, such as nicotinic acetylcholine (nAchR), type-3 somatostatin, and GABA-A receptors (Jamin *et al.*, 2005; Epand, 2006; Fantini & Barrantes, 2013).

We recently reported identification of a CARC domain in TRKB and showed that mutation in the CARC domain interferes with plasticity-related BDNF signaling (Casarotto *et al.*, 2021). The aim of the present study was to evaluate the incidence of cholesterol-interacting motifs (CRAC and CARC) in the RTK family. Given the promiscuous nature of CRAC motifs, we focused on the RTK transmembrane region (TMR; transmembrane domain plus the 5-amino acid flanking residues on both N- and C-terminal sides), where such interaction has a higher probability of occurrence. Transmembrane domains are crucial for proper positioning in the lipid bilayer of biological membranes (Fantini & Barrantes, 2013). Interaction of transmembrane domains of embedded integral proteins with the lipid component of the bilayer provides a diffusion barrier and encloses the environment to maintain electrochemical properties (Hunte, 2005). Upon identification of CRAC/CARC motifs in the TMR of many members of the RTK family, we focused on TRKB and assessed its percentage of identity across species. We then investigated the interaction of this motif with cholesterol, by molecular dynamic simulations. Then, we expanded previous findings about the mechanisms behind the cholesterol effect on BDNF-induced TRKB activity (Casarotto *et al.*, 2021) by assaying the binding of biotinylated BDNF to immobilized TRKB.

## Methods

### Data mining

For data mining, we used 144 manually curated inputs of RTK family (code 2.7.10.1) from the UniProt database (The UniProt Consortium, 2017). The canonical primary structure of TMR (transmembrane domain and the flanking 5 amino acid residues, from N- and C-terminal) of RTK from each target of human (52 proteins), mouse (51 proteins), zebrafish (14 proteins), fruit fly (12 proteins), and nematode *C. elegans* (15 proteins) databases were extracted. The TMR FASTA sequences for each protein were manually screened for the presence of cholesterol recognition alignment consensus, both CRAC and CARC (Fantini & Barrantes, 2013; Fantini *et al.*, 2016). We then searched for putative pathogenic mutations in human proteins using SwissVar, ClinVar, and COSMIC databases (Mottaz *et al.*, 2010; Landrum *et al.*, 2018; Tate *et al.*, 2019).

### Percentage of identity of TRKB (and TMR) across species

The Percentage Identity (PI) of TRKB.TMR (and full-length sequences) among several species, including *D. rerio* (zebrafish), *G. gallus* (chicken), *C. familiaris* (dog), *R. norvegicus* (rat), *M. musculus* (mouse), *P. troglodytes* (chimpanzee), and *H. sapiens* (human), was determined using the align tool in UniProt database (The UniProt Consortium, 2017). The correlation between PI in full-length TRKB and TRKB.TMR among the different species was determined by Spearman’s test (GraphPad Prism v.6.07). The PI trees for full TRKB and TMR.TRKB were obtained using the tree tool (Neighbour Joining, BLOSUM62) in Jalview v.2.0 software (Waterhouse *et* al., 2009).

### Molecular dynamics simulations

The 3-dimensional (3D) structure of the TMR of TRKB (residues 423-460) was generated using the FMAP (Folding of Membrane-Associated Peptides) server (Lomize *et al.*, 2018). The server predicted that the residues V432-A456 form an α-helical transmembrane segment; the remaining sequence is unstructured. This structure was used as an atomistic model for the TRKB. For coarse-grained simulations, these atomistic structures were used as the basis. For this purpose, all protein-membrane systems were constructed using the CHARMM-GUI Martini Maker (Hsu *et al.*, 2017), where the TRKB was separately embedded in a bilayer (512 lipids) composed of 60 mol% palmitoyl-oleoyl-phosphatidylcholine (POPC) and 40 mol% cholesterol. Each system was solvated with 6038 water beads (approximately 50 water molecules per lipid). Sodium and chloride ions were added to reach 0.15 M salt concentration and to neutralize the system.

All simulations were performed using Gromacs 5.1.4 (Abraham *et al.*, 2015) employing the non-polarizable Martini 2.2 force field for the protein (de Jong *et al.*, 2013) and lipids (Arnarez *et al.*, 2015). The simulations were performed using the “New-RF” parameters (de Jong *et al.*, 2016). For electrostatics, the reaction field method was used with a cutoff of 1.1 nm. Lennard-Jones interactions were cut off at 1.1 nm. The potential shift modifier was applied to non-bonded interactions together with buffered Verlet lists (Páll & Hess, 2013). The equations of motion were integrated using the leap-frog algorithm with a 25-fs time step. The simulations were performed at 310 K in the NpT ensemble at a pressure of 1 bar. The protein, the membrane, and the solvent (water and 0.15 M NaCl) were coupled to separate heat baths with a time constant of 1.0 ps using the V-rescale thermostat (Bussi *et al.*, 2007). Pressure was controlled semi-isotropically using the Parrinello-Rahman barostat (Parrinello & Rahman, 1981) with a time constant of 12 ps and a compressibility of 3×10^−4^ bar^−1^ in the xy-plane (membrane plane). The systems were first equilibrated with the protein backbone atoms restrained. In the production stage, each system (TRKB.wt and TRKB.mut) was simulated for 5 μs through 9 independent repeats.

All analyses were performed using the Gromacs software package and in-house scripts, using only the last 4 μs of the simulations. The data presented (figure 3) is the average occupancy of cholesterol (occupancy) that represents the average number of cholesterol molecules within 0.6 nm of the alpha carbon (MARTINI backbone bead) at residue positions 427 and 433. The occupancy values are produced by dividing the total number of contacts by the number of frames in each trajectory.

### Cell cultures and BDNF binding assay

HEK293T cells were transfected to overexpress TRKB. The cells were maintained at 5% CO2, 37°C in Dulbecco’s Modified Eagle’s Medium (DMEM, containing 10% fetal calf serum, 1% penicillin/streptomycin, 1% L-glutamine). The cells were lysed, and the lysate was submitted to the BDNF binding assay.

The BDNF binding to TRKB was performed in white 96-well plates (Das *et al.*, 2015; Baeza-Raja *et al.*, 2016; Casarotto *et al.*, 2021). Briefly, the plates were precoated with anti-TRKB antibody (1:1000, R&D Systems, #AF1494) in carbonate buffer (pH 9.8) overnight at 4°C, followed by blocking with 3% BSA in PBS buffer (2 h at RT). The samples (120ug of total protein) were added and incubated overnight at 4°C under agitation. The plates were washed 3x with PBS buffer, and a mixture of biotinylated BDNF (bBDNF, 0, 0.1, 1 or 10ng/ml, Alomone Labs, #B-250-B) and cholesterol (0, 20, 50 or 100μM) was added for 1h at room temperature, followed by washing with PBS. The luminescence was determined via HRP-conjugated streptavidin (1:10000, 1h, RT, Thermo-Fisher, #21126) activity reaction with ECL by a plate reader. The luminescence signal from blank wells (containing all the reagents but the sample lystates, substituted by the blocking buffer) was used as background. The specific signal was then calculated by subtracting the values of blank wells from the values of the samples with matched concentration of the biotinylated ligand. The signal was normalized by the bBDNF at 10ng/ml under no added cholesterol (0 μM).

### Statistical analysis

The data of the present manuscript was analyzed by Student t test for MD simulations or two-way ANOVA for BDNF binding. The data used in the present study is available in FigShare under CC-BY license (DOI:10.6084/m9.figshare.7935980).

## Results

### Data mining

The presence of CRAC motifs within the TMDs of RTK family members was found throughout all the species analyzed (human, 11 of 52 proteins; mouse, 10 of 51 proteins; zebrafish, 2 of 14 proteins; fruit fly, 2 of 12 proteins; and C. elegans, 2 of 15 proteins, figure 1). However, the presence of CARC motifs in the RTK family was observed only in vertebrates, with 3 in human, 3 in mouse, and 2 in zebrafish RTK. None of the proteins analyzed was found to carry CRAC and CARC motifs simultaneously. The ClinVar and COSMIC databases indicated five proteins (Table 1) consisting of 8 mutations in the CRAC/CARC domains associated with central nervous system or endocrine disorders or cancer.

**Table 1.**
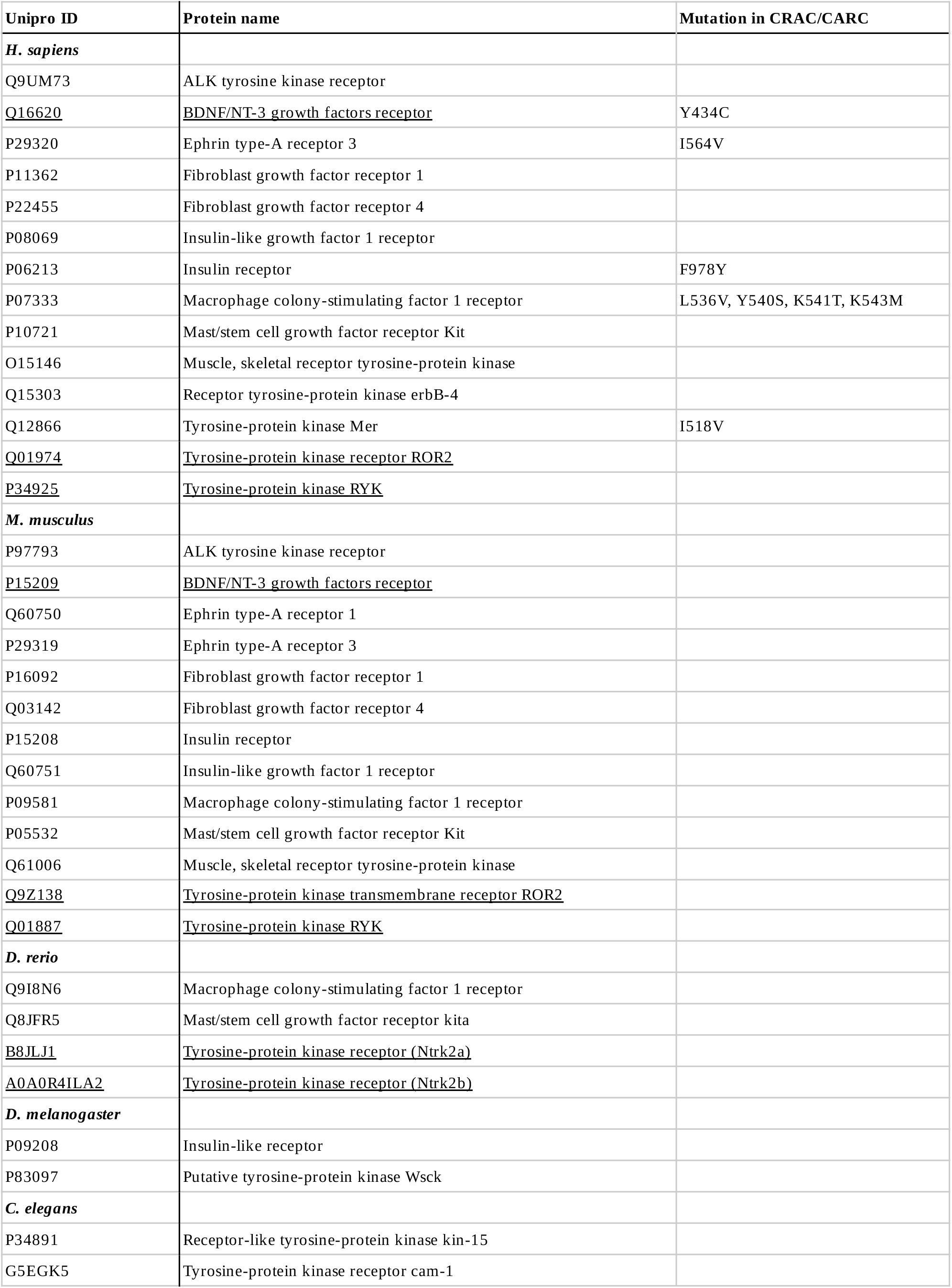
CRAC and CARC(underlined)-positive proteins.

| Unipro ID | Protein name | Mutation in CRAC/CARC |
| --- | --- | --- |
| *H. sapiens* |  |  |
| Q9UM73 | ALK tyrosine kinase receptor |  |
| <u>Q16620</u> | <u>BDNF/NT-3 growth factors receptor</u> | Y434C |
| P29320 | Ephrin type-A receptor 3 | I564V |
| P11362 | Fibroblast growth factor receptor 1 |  |
| P22455 | Fibroblast growth factor receptor 4 |  |
| P08069 | Insulin-like growth factor 1 receptor |  |
| P06213 | Insulin receptor | F978Y |
| P07333 | Macrophage colony-stimulating factor 1 receptor | L536V, Y540S, K541T, K543M |
| P10721 | Mast/stem cell growth factor receptor Kit |  |
| O15146 | Muscle, skeletal receptor tyrosine-protein kinase |  |
| Q15303 | Receptor tyrosine-protein kinase erbB-4 |  |
| Q12866 | Tyrosine-protein kinase Mer | I518V |
| <u>Q01974</u> | <u>Tyrosine-protein kinase receptor ROR2</u> |  |
| <u>P34925</u> | <u>Tyrosine-protein kinase RYK</u> |  |
| *M. musculus* |  |  |
| P97793 | ALK tyrosine kinase receptor |  |
| <u>P15209</u> | <u>BDNF/NT-3 growth factors receptor</u> |  |
| Q60750 | Ephrin type-A receptor 1 |  |
| P29319 | Ephrin type-A receptor 3 |  |
| P16092 | Fibroblast growth factor receptor 1 |  |
| Q03142 | Fibroblast growth factor receptor 4 |  |
| P15208 | Insulin receptor |  |
| Q60751 | Insulin-like growth factor 1 receptor |  |
| P09581 | Macrophage colony-stimulating factor 1 receptor |  |
| P05532 | Mast/stem cell growth factor receptor Kit |  |
| Q61006 | Muscle, skeletal receptor tyrosine-protein kinase |  |
| <u>Q9Z138</u> | <u>Tyrosine-protein kinase transmembrane receptor ROR2</u> |  |
| <u>Q01887</u> | <u>Tyrosine-protein kinase RYK</u> |  |
| *D. rerio* |  |  |
| Q9I8N6 | Macrophage colony-stimulating factor 1 receptor |  |
| Q8JFR5 | Mast/stem cell growth factor receptor kita |  |
| <u>B8JLJ1</u> | <u>Tyrosine-protein kinase receptor (Ntrk2a)</u> |  |
| <u>A0A0R4ILA2</u> | <u>Tyrosine-protein kinase receptor (Ntrk2b)</u> |  |
| *D. melanogaster* |  |  |
| P09208 | Insulin-like receptor |  |
| P83097 | Putative tyrosine-protein kinase Wsck |  |
| *C. elegans* |  |  |
| P34891 | Receptor-like tyrosine-protein kinase kin-15 |  |
| G5EGK5 | Tyrosine-protein kinase receptor cam-1 |  |

**Figure 1.**
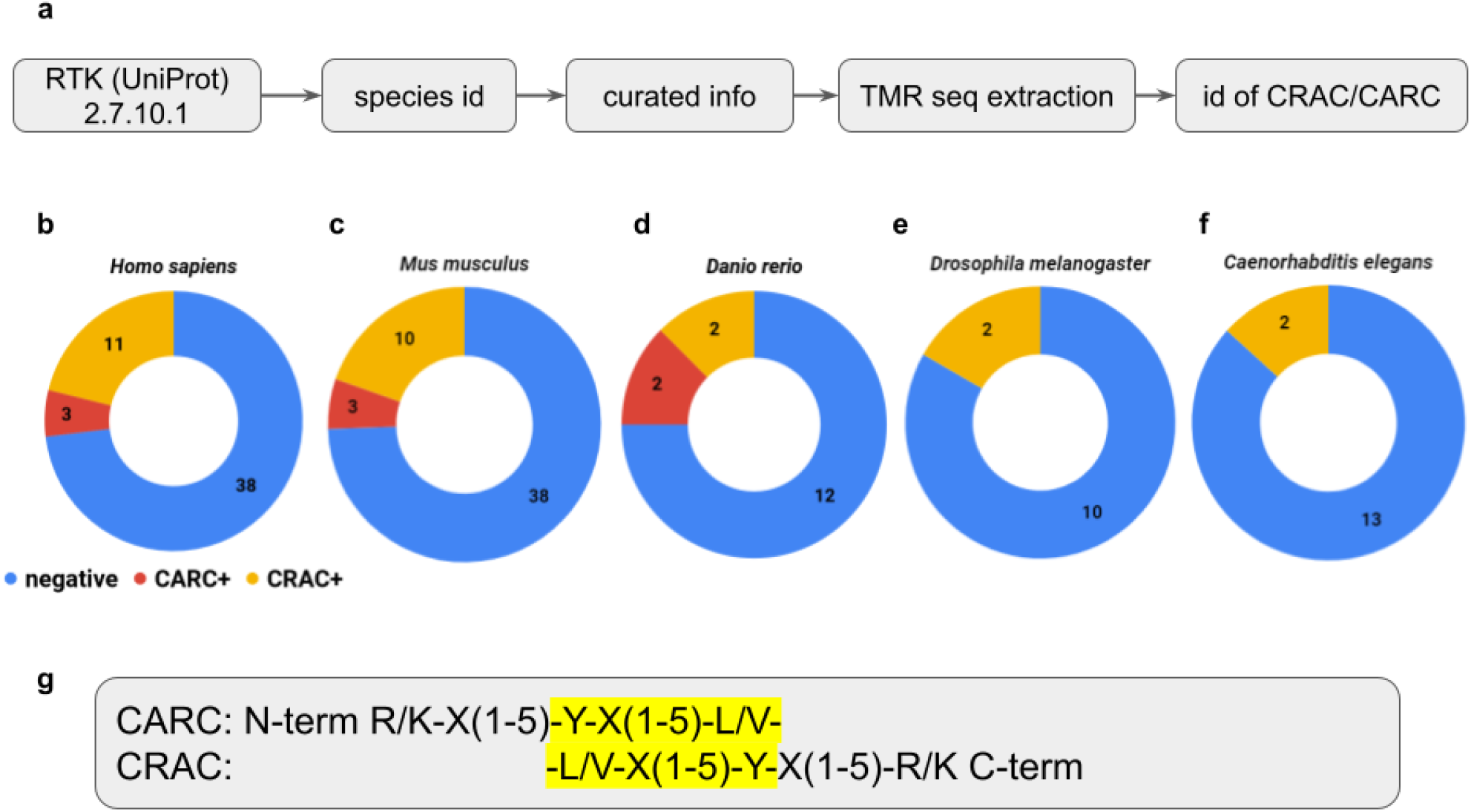
**(a)** Workflow of data mining. The TMR (transmembrane sequence +5 amino acid residues from each N- and C-terminal sides) of curated entries found in UniProt using the code for tyrosine kinase receptor family (2.7.10.1) were blasted against the library of cholesterol recognition and alignment consensus combinations. The incidence of CRAC and CARC motifs in **(b)** human, **(c)** mouse, **(d)** zebrafish, **(e)** fruit fly, and **(f)** *C. elegans* TMR of RTK family members. **(g)** Library of CRAC and CARC sequences (all combinations used can be found in the stored data). The yellow highlighted region of the sequences must be embedded into the cell membrane.

The full list of proteins positive to CRAC/CARC is found in table 1 and the full list of proteins examined from each species can be found in the deposited data.

### Percentage of identity of TRKB (and TMR) across species

We previously found that TRKB is the only member of the TRK-subfamily of RTKs that possesses a CARC domain (Casarotto *et al.*, 2021). The CARC motif in TRKB was conserved across species: human and mouse, REHLSVYAVVV; zebrafish, RVAVYIVV. The identity between TRKB.TMR sequences of several species (human, chimpanzee, mouse, rat, dog, chicken, and zebrafish; table 2) was examined using UniProt (The UniProt Consortium, 2017). Over 90% PI was found in both TRKB.TMR and full-length TRKB of human, chimpanzee, mouse, rat, and dog. The PI results of paired comparison between the species analyzed are organized in figure 2, where the green gradient highlights the percentage similarity between the sequences. For comparative purposes, we also determined the PI of full-length TRKB among these species. The PI results of paired comparisons are also organized in figure 2, with red gradient highlighting similarity. Spearman’s test indicated a significant correlation between the full-length TRKB and TRKB.TMR [R^2^=0.8530, 99% confidence interval, CI=0.8135-0.9698; p<0.0001].

**Table 2.** CARC-containing sequences (red) in TRKB.TMR among vertebrate species.

| UniProt | Species | TMR sequence |
| --- | --- | --- |
| Q16620 | *H. sapiens* | TGREHLSVYAVVVIASVVGFCLLVMLFLLKLARH |
| A0A2J8MRP9 | *P. troglodites* | TGREHLSVYAVVVIASVVGFCLLVMLFLLKLARH |
| P15209 | *M. musculus* | SNREHLSVYAVVVIASVVGFCLLVMLLLLKLARH |
| Q63604 | *R. norvegicus* | TNREHLSVYAVVVIASVVGFCLLVMLLLLKLARH |
| E2RKA1 | *C. familiaris* | SGREHLSVYAVVVIASVVGFCLLVMLFLLKLARH |
| Q91987 | *G. gallus* | ENEDSITVYVVVGIAALVCTGLVIMLIILKFGRH |
| A0A0R4ILA2 | *D. rerio* | PLEDRVAVYIVVGIAGVALTGCILMLVFLKYGRS |

**Figure 2.**
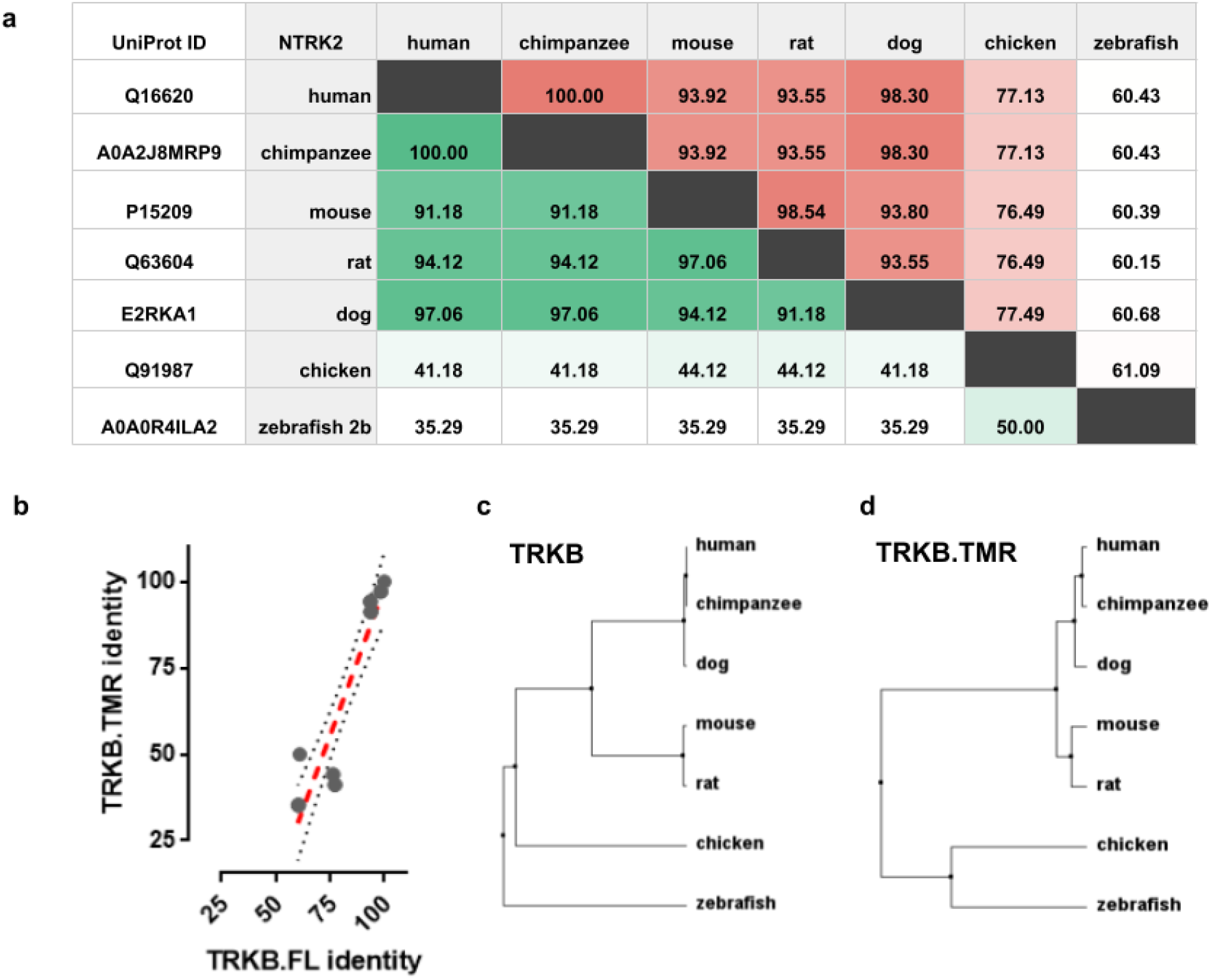
**(a)** Percent identity (PI) for full length (red) and TMR (green) of TRKB. The amino acid sequence of full length and TMR of TRKB (NTRK2) from different species were verified for PI in the UniProt database. **(b)** Correlation between the PI in full length and TMR of TRKB [Spearman’s test. R2= 0.8530; 99%CI= 0.7564 to 0.9775; p<0.0001]; dashed line= 99% confidence band. PI tree of **(c)** full length and **(d)** TMR of TRKB.

**Figure 3.**
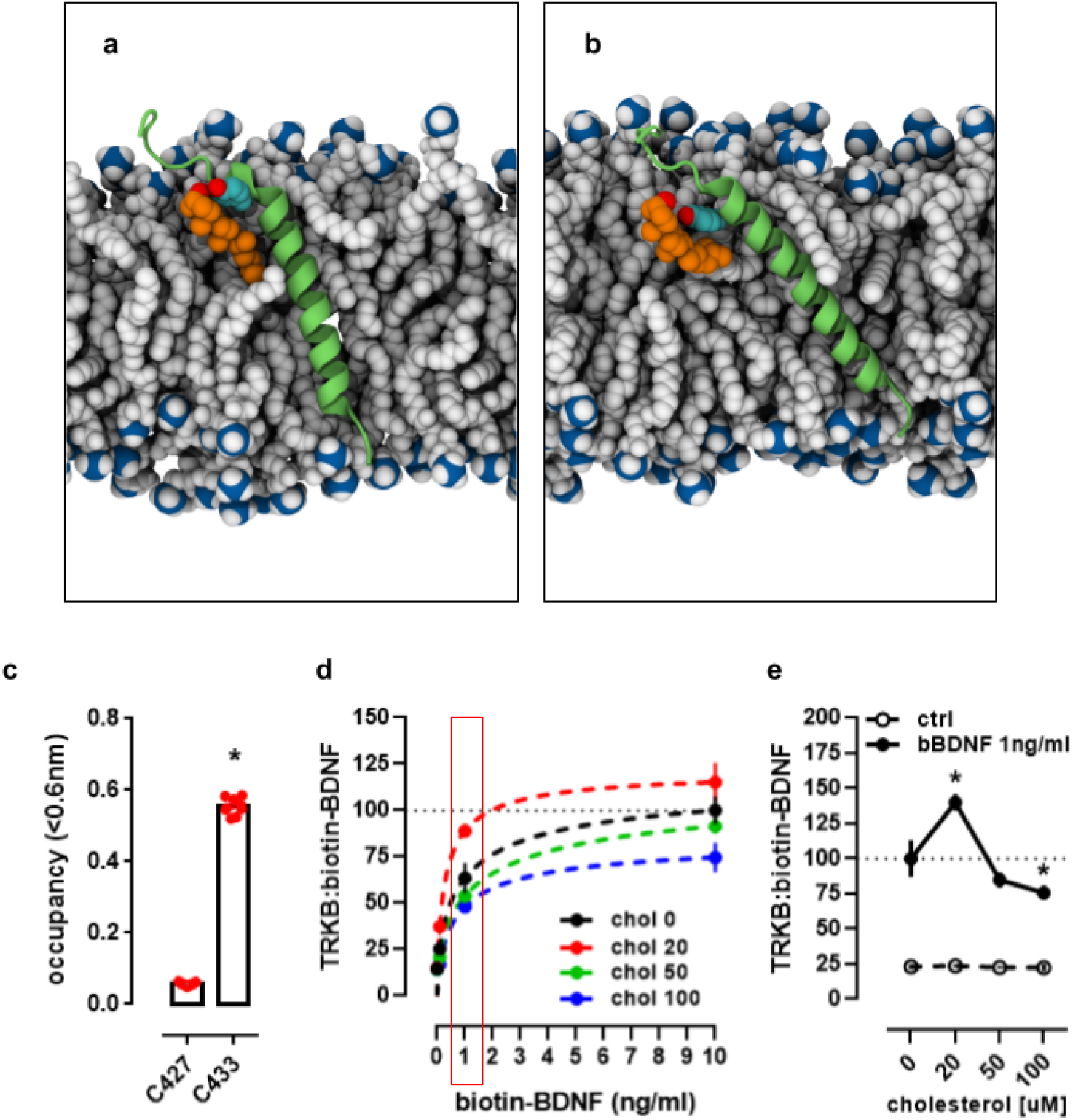
TRKB and cholesterol interaction. **(a,b)** Snapshots of the interaction between TRKB and cholesterol at the TMR (see supplement video) indicating that cholesterol interacts with the OH- group (red) in Y433 (blue) in TRKB.TMR (green). **(c)** The interaction between the TRKB CARC motif and cholesterol happens preferentially at the C-alpha in Y433 residues, as indicated by coarse-grained molecular dynamics simulations. **(d)** The binding of biotinylated BDNF (bBDNF) and TRKB is modulated by cholesterol in a bell-shaped fashion. Red rectangle area is expanded in **(e)** for the comparison between BDNF 1ng/ml (black circles) with BDNF 0 (ctrl, open circles). Data expressed as Mean/SEM of **(c)** cholesterol occupancy; or from binding normalized by **(d)** chol 0/BDNF10ng/ml or **(e)** chol 0/BDNF 1ng/ml. ∗p<0.05 from control (BDNF 0 or ctrl, C427).

### Molecular dynamics simulations

To study the interaction between cholesterol and TRKB TMR, we performed coarse-grained simulations embedded in a 40 mol% cholesterol bilayer, snapshots of the trajectories can be seen in figure 3a,b and in supplement video. Analysis of the trajectories in terms of average cholesterol occupancy (see Methods) revealed a higher occupancy of cholesterol molecules in the Y433 compared to R427 of the TRKB CARC domain [t(16)=62.46, p<0.0001], figure 3c.

### BDNF binding assay

As found in figure 3d, the binding of bBDNF in immobilized TRKB is altered by cholesterol concentration. The two-way ANOVA indicated an interaction between the cholesterol and BDNF binding [F(9,80)= 2.987, p= 0.0041]. Specifically, cholesterol exerts a bell-shaped effect in BDNF binding to TRKB, where lower concentrations of cholesterol (20μM) facilitate, while higher amounts of cholesterol (50 or 100μM) compromise bBDNF detection in immobilized TRKB, as evidenced by the curve for bBDNF at 1ng/ml under different concentrations of cholesterol [F(3,40)= 11.50, p< 0.0001], figure 3e.

## Discussion

In the present study, we evaluated the incidence of cholesterol-recognition motifs in the RTK family from mouse, human, zebrafish, fruit fly, and the nematode *C. elegans*. We found that while the *bona fide* CRAC, located in the C-terminal portion of the transmembrane domain (Fantini & Barrantes, 2013), is found in all species analyzed, its inverted version (CARC) was observed only in vertebrates. Furthermore, we found that TRKB, a tyrosine-kinase receptor crucial for neuronal plasticity, contains a CARC sequence in the TMR that is conserved across different species. However, CRAC/CARC motifs were absent in the TMR of other members of the TRK subfamily, such as TRKA and TRKC.

Cholesterol can interact with membrane proteins in several ways. One of its most prominent effects involves a direct post-translational modification on members of the Hedgehog pathway described in *Drosophila sp*. (Jeong & McMahon, 2002; Wendler *et al.*, 2006). In this model organism, cholesterol also regulates membrane depolarization through transient receptor potential (TRP) channels (Peters, 2017) and serves as a precursor for ecdysteroids, which in turn control several steps of the fly development (Niwa & Niwa, 2014). In nematodes such as C. elegans, cholesterol is only obtained from diet, although these worms can modify the basic steroid structure into derivatives (Kurzchalia & Ward, 2003). In both organisms, cholesterol appears to play a major role as a signaling molecule with post-translational modifications of proteins as the main mechanism (Mann & Beachy, 2000).

In vertebrates, although neurons synthesize the absolute minimum of required cholesterol, glial production and release of lipoproteins supply neuronal demand during development and in adulthood (Mauch *et al.*, 2001). In particular, apolipoprotein E (APOE) is synthesized primarily by astrocytes and glial cells (Boyles *et al.*, 1985; Pfrieger & Ungerer, 2011). Glia-derived cholesterol stimulates synapse formation and synaptic efficacy (Pfrieger, 2003a, 2003b). In the presynaptic plasma membrane, cholesterol-rich lipid rafts are necessary for SNARE-dependent exocytosis of vesicles with high cholesterol content. At the postsynaptic level, such rafts organize the disposition of receptors, protein scaffolds, and signaling cascades (Pfrieger, 2003a, 2003b). Importantly, cholesterol removal from neuronal cultures impairs exocytosis of synaptic vesicles (Linetti *et al.*, 2010), *s*ynaptic transmission (Goritz *et al.*, 2005), and neuronal viability (Michikawa & Yanagisawa, 1999). In addition, cholesterol induces clustering of AMPA receptors and hinders NMDA-induced long-term potentiation in the hippocampus (Frank *et al.*, 2008; Martín *et al*., 2014).

Two consensus motifs with predictive value for cholesterol interaction with proteins have been defined through *in silico* methods (Di Scala *et al.*, 2017), CRAC and CARC (Li & Papadopoulos, 1998; Baier *et al.*, 2011). The non-covalent binding of cholesterol to such motifs has been the focus of various recent studies. For example, cholesterol modulates docking of NMDA receptors into lipid rafts (Korinek *et al.*, 2015) and regulates the function of vanilloid receptors TRPV1, a member of the TRP family (Jansson *et al.*, 2013), thus interfering in synaptic plasticity. Increased cholesterol concentration enhances the plasticity and flexibility of 5HT1a dimers and adrenergic receptors (Prasanna *et al.*, 2014, 2016). Given the opposed dispositions of CARC and CRAC motifs, it is possible to assume the co-existence of both in the same transmembrane domain and their potential interaction with two cholesterol molecules in a tail-to-tail configuration (Di Scala *et al.*, 2017). However, none of the analyzed TMR of RTK family members in the present study displayed co-existing CARC and CRAC motifs.

Interestingly, we only observed the occurrence of CARC motifs in the zebrafish, mouse, and human RTK family. Only three vertebrate RTKs: NTRK2 (TRKB), ROR2 and RYK, were found to possess a CARC domain within the TMR. ROR2 is a member of the ROR family, closely related to the TRK family, and plays a distinct role in bone morphogenesis, through a signaling cascade not yet fully described but engaging 14-3-3β scaffolding proteins (Liu *et al.*, 2007). RYK receptor is an atypical member of the RTK family as it lacks tyrosine kinase activity while containing the related domain (Hovens *et al.*, 1992). In mammalian cells RYK serves as a co-receptor with Frizzled for Wnt ligands mediating neurite outgrowth (Lu *et al.*, 2004).

TRKB plays a crucial role in several aspects of neuronal plasticity (Park & Poo, 2013). The activation of this receptor is associated with the reopening of the visual critical period (Maya Vetencourt *et al.*, 2008) and the formation, retention, and recall of memory (Karpova *et al.*, 2011; Bekinschtein *et al.*, 2014). TRKB.TMR is highly conserved among vertebrates, similar to full-length TRKB. Although correlated, the identity of full-length TRKB and TRKB.TMR are not comparable. Given the large difference in the number of residues between these two sequences, each residue change in the TMR exerts a higher impact than on full-length TRKB in the overall identity between the species analyzed. However, the TRKB.TMR CARC sequence from chicken differs in the juxtamembrane residues from the other species compared here (table 2). The following two scenarios are plausible. The role of R/K (charged, basic residues) is fulfilled by glutamate (E), which is also charged at pH 7, although negatively; or by asparagine (N), which is not charged but carries a basic amino group. Additionally, for the second possibility it is also necessary to relax the proposed “5-residue rule” between the Y and the juxtamembrane residue (Fantini *et al.*, 2019) since N is located 6 residues apart from the central Y. Nonetheless, our MD simulations indicate that cholesterol interacts with TRKB mainly through the central Y residue, as the interaction values between the C-alpha in Y433 residue (C433) are 10-fold higher than those between cholesterol and the C427, suggesting that chicken TRKB might still be able to interact with cholesterol. Moreover, the cholesterol interaction with TRKB, measured by microscale thermophoresis, is completely lost in the single mutant Y433F, reinforcing the central role for Y in this interaction (Casarotto *et al.*, 2021).

TRKB is found in lipid rafts only upon activation by BDNF (Suzuki *et al.*, 2004). Interestingly, when cholesterol is sequestered, TRKB translocation to lipid rafts is impaired, and BDNF-dependent potentiation is prevented (Suzuki *et al.*, 2004). However, loss of cholesterol in hippocampal cultures is associated with increased baseline activity of TRKB (Martin *et al.*, 2008). These opposite outcomes might be due to a differential modulation exerted by cholesterol, depending on the challenge to TRKB receptor (basal vs BDNF-stimulated), cell type or origin, and stage of differentiation. Another possibility is that cholesterol affects TRKB activity in a bell-shaped manner, where higher and lower cholesterol concentrations impede instead of promote TRKB phosphorylation. In fact, the decrease of cholesterol levels by beta-cyclodextrin was found to differentially modulate neurite growth of hippocampal and cortical cultured neurons (Ko *et al.*, 2005). In hippocampal cells, the decrease of cholesterol levels induced an increase in neurite length and number, while no effect was observed in cortical cells. Interestingly, cultures of hippocampal cells revealed higher levels of cholesterol than the cortical counterparts (Ko *et al.*, 2005).

Altogether, it is a provocative idea to consider TRKB as a “sensor” of cholesterol levels in the cell membrane via CARC. Thus, TRKB would trigger synaptic maturation or neurite growth only if the cholesterol levels are ideal for such changes, *i.e*. cholesterol concentrations must be within a ‘Goldilocks’ zone. In fact, our previous data indicates that cholesterol indeed modulates BDNF-induced TRKB activation in a bell-shaped fashion (Casarotto *et al.*, 2021). Under low amounts of supplemented cholesterol, the activation of TRKB by BDNF is facilitated, while at higher concentrations (around 50 μM) this increment is lost, and even compromised under higher amounts of cholesterol (around 100 μM), and inhibiting cholesterol synthesis also compromised BDNF effect (Suzuki *et al.*, 2004; Casarotto *et al.*, 2021). In line with this evidence, here we observed the same pattern over BDNF binding to TRKB. TRKB dimerization is heavily influenced by cholesterol, mainly through changes in the membrane thickness, which changes the orientation of TRKB dimers between signaling competent and incompetend conformations (Casarotto *et al.*, 2021), and the present data provides additional evidence that such changes may also influence BDNF interaction with TRKB.

The insights from the present study could serve as a primary step for experimental testing on the impact of mutations in CRAC/CARC motifs in the TMR of the RTK family. However, we are limited by only considering the role of CRAC/CARC motifs in the TMR of RTKs. Given the promiscuous properties of these motifs, it is plausible to assume multiple false positive CRAC/CARCs in proteins, making data mining and putative *in silico* or *in vitro* analysis difficult to perform. Therefore, more studies focused on refining the algorithms for detecting these motifs are necessary.

## Acknowledgements

The authors thank Sulo Kolehmainen and Seija Lågas for their technical assistance. This study was funded by grants from European Research Council (#322742), EU Joint Programme -Neurodegenerative Disease Research (JPND) CircProt (#301225 and #643417), Sigrid Jusélius Foundation, Jane and Aatos Erkko Foundation, and the Academy of Finland (#294710 and #307416). None of the funders had a role in the data acquisition, analysis or manuscript preparation.

## Contributions and conflict of interest

CC, MF, PC, CB conducted the data mining and binding experiments. MG, GE and TR conducted the molecular dynamics experiments. CC, MG, GE, PC, CB and MF wrote the first draft of the manuscript, edited by TR, IV and EC. EC is a shareholder of Herantis Pharma, and received lecture fees from Janssen-Cilag. All other authors declare no competing interests.

## References

Abraham, M.J., Murtola, T., Schulz, R., Páll, S., Smith, J.C., Hess, B., & Lindahl, E. (2015) GROMACS: High performance molecular simulations through multi-level parallelism from laptops to supercomputers. SoftwareX, 1-2, 19–25.

Arnarez, C., Uusitalo, J.J., Masman, M.F., Ingólfsson, H.I., de Jong, D.H., Melo, M.N., Periole, X., de Vries, A.H., & Marrink, S.J. (2015) Dry Martini, a coarse-grained force field for lipid membrane simulations with implicit solvent. J. Chem. Theory Comput., 11, 260–275.

Baeza-Raja, B., Sachs, B.D., Li, P., Christian, F., Vagena, E., Davalos, D., Le Moan, N., Ryu, J.K., Sikorski, S.L., Chan, J.P., Scadeng, M., Taylor, S.S., Houslay, M.D., Baillie, G.S., Saltiel, A.R., Olefsky, J.M., & Akassoglou, K. (2016) p75 Neurotrophin Receptor Regulates Energy Balance in Obesity. Cell Rep., 14, 255–268.

Baier, C.J., Fantini, J., & Barrantes, F.J. (2011) Disclosure of cholesterol recognition motifs in transmembrane domains of the human nicotinic acetylcholine receptor. Sci. Rep., 1, 69.

Bekinschtein, P., Cammarota, M., & Medina, J.H. (2014) BDNF and memory processing. Neuropharmacology, 76 Pt C, 677–683.

Boyles, J.K., Pitas, R.E., Wilson, E., Mahley, R.W., & Taylor, J.M. (1985) Apolipoprotein E associated with astrocytic glia of the central nervous system and with nonmyelinating glia of the peripheral nervous system. J. Clin. Invest., 76, 1501–1513.

Bussi, G., Donadio, D., & Parrinello, M. (2007) Canonical sampling through velocity rescaling. J. Chem. Phys., 126, 014101.

Casarotto, P.C., Girych, M., Fred, S.M., Moliner, R., Enkavi, G., Biojone, C., Cannarozzo, C., Brunello, C.A., Steinzeig, A., Winkel, F., Patil, S., Vestring, S., Serchov, T., Kovaleva, V., Diniz, C.R., Laukkanen, L., Cardon, I., Antila, H., Rog, T., Saarma, M., Bramham, C.R., Normann, C., Lauri, S.E., Vattulainen, I., & Castrén, E. (2021) Antidepressant drugs act by directly binding to TRKB neurotrophin receptors. Cell, in press.

Das, I., Krzyzosiak, A., Schneider, K., Wrabetz, L., D’Antonio, M., Barry, N., Sigurdardottir, A., & Bertolotti, A. (2015) Preventing proteostasis diseases by selective inhibition of a phosphatase regulatory subunit. Science, 348, 239–242.

de Chaves, E.I., Rusiñol, A.E., Vance, D.E., Campenot, R.B., & Vance, J.E. (1997) Role of lipoproteins in the delivery of lipids to axons during axonal regeneration. J. Biol. Chem., 272, 30766–30773.

de Jong, D.H., Baoukina, S., Ingólfsson, H.I., & Marrink, S.J. (2016) Martini straight: Boosting performance using a shorter cutoff and GPUs. Comput. Phys. Commun., 199, 1–7.

de Jong, D.H., Singh, G., Bennett, W.F.D., Arnarez, C., Wassenaar, T.A., Schäfer, L.V., Periole, X., Tieleman, D.P., & Marrink, S.J. (2013) Improved Parameters for the Martini Coarse-Grained Protein Force Field. J. Chem. Theory Comput., 9, 687–697.

Dekkers, M.P.J., Nikoletopoulou, V., & Barde, Y.-A. (2013) Cell biology in neuroscience: Death of developing neurons: new insights and implications for connectivity. J. Cell Biol., 203, 385–393.

Dietschy, J.M. & Turley, S.D. (2004) Thematic review series: Brain Lipids. Cholesterol metabolism in the central nervous system during early development and in the mature animal. J. Lipid Res., 45, 1375–1397.

Di Scala, C., Baier, C.J., Evans, L.S., Williamson, P.T.F., Fantini, J., & Barrantes, F.J. (2017) Chapter One - Relevance of CARC and CRAC Cholesterol-Recognition Motifs in the Nicotinic Acetylcholine Receptor and Other Membrane-Bound Receptors. In Levitan, I. (ed), Current Topics in Membranes, Sterol Regulation of Ion Channels. Academic Press, pp. 3–23.

Elkins, M.R., Sergeyev, I.V., & Hong, M. (2018) Determining Cholesterol Binding to Membrane Proteins by Cholesterol 13C Labeling in Yeast and Dynamic Nuclear Polarization NMR. J. Am. Chem. Soc., 140, 15437–15449.

Epand, R.M. (2006) Cholesterol and the interaction of proteins with membrane domains. Prog. Lipid Res., 45, 279–294.

Fantini, J. & Barrantes, F.J. (2013) How cholesterol interacts with membrane proteins: an exploration of cholesterol-binding sites including CRAC, CARC, and tilted domains. Front. Physiol., 4, 31.

Fantini, J., Di Scala, C., Evans, L.S., Williamson, P.T.F., & Barrantes, F.J. (2016) A mirror code for protein-cholesterol interactions in the two leaflets of biological membranes. Sci. Rep., 6, 21907.

Fantini, J., Epand, R.M., & Barrantes, F.J. (2019) Cholesterol-Recognition Motifs in Membrane Proteins. Adv. Exp. Med. Biol., 1135, 3–25.

Frank, C., Rufini, S., Tancredi, V., Forcina, R., Grossi, D., & D’Arcangelo, G. (2008) Cholesterol depletion inhibits synaptic transmission and synaptic plasticity in rat hippocampus. Exp. Neurol., 212, 407–414.

Goritz, C., Mauch, D.H., & Pfrieger, F.W. (2005) Multiple mechanisms mediate cholesterol-induced synaptogenesis in a CNS neuron. Mol. Cell. Neurosci., 29, 190–201.

Hanson, M.A., Cherezov, V., Griffith, M.T., Roth, C.B., Jaakola, V.-P., Chien, E.Y.T., Velasquez, J., Kuhn, P., & Stevens, R.C. (2008) A specific cholesterol binding site is established by the 2.8 A structure of the human beta2-adrenergic receptor. Structure, 16, 897–905.

Harris, J.S., Epps, D.E., Davio, S.R., & Kézdy, F.J. (1995) Evidence for transbilayer, tail-to-tail cholesterol dimers in dipalmitoylglycerophosphocholine liposomes. Biochemistry, 34, 3851–3857.

Hovens, C.M., Stacker, S.A., Andres, A.C., Harpur, A.G., Ziemiecki, A., & Wilks, A.F. (1992) RYK, a receptor tyrosine kinase-related molecule with unusual kinase domain motifs. Proc. Natl. Acad. Sci. U. S. A., 89, 11818–11822.

Hsu, P.-C., Bruininks, B.M.H., Jefferies, D., Cesar Telles de Souza, P., Lee, J., Patel, D.S., Marrink, S.J., Qi, Y., Khalid, S., & Im, W. (2017) CHARMM-GUI Martini Maker for modeling and simulation of complex bacterial membranes with lipopolysaccharides. J. Comput. Chem., 38, 2354–2363.

Huang, E.J. & Reichardt, L.F. (2001) Neurotrophins: roles in neuronal development and function. Annu. Rev. Neurosci., 24, 677–736.

Hunte, C. (2005) Specific protein-lipid interactions in membrane proteins. Biochem. Soc. Trans., 33, 938–942.

Jamin, N., Neumann, J.-M., Ostuni, M.A., Vu, T.K.N., Yao, Z.-X., Murail, S., Robert, J.-C., Giatzakis, C., Papadopoulos, V., & Lacapère, J.-J. (2005) Characterization of the Cholesterol Recognition Amino Acid Consensus Sequence of the Peripheral-Type Benzodiazepine Receptor. Mol. Endocrinol., 19, 588–594.

Jansson, E.T., Trkulja, C.L., Ahemaiti, A., Millingen, M., Jeffries, G.D., Jardemark, K., & Orwar, O. (2013) Effect of cholesterol depletion on the pore dilation of TRPV1. Mol. Pain, 9, 1.

Jeong, J. & McMahon, A.P. (2002) Cholesterol modification of Hedgehog family proteins. J. Clin. Invest., 110, 591–596.

Karpova, N.N., Pickenhagen, A., Lindholm, J., Tiraboschi, E., Kulesskaya, N., Agústsdóttir, A., Antila, H., Popova, D., Akamine, Y., Bahi, A., Sullivan, R., Hen, R., Drew, L.J., & Castrén, E. (2011) Fear erasure in mice requires synergy between antidepressant drugs and extinction training. Science, 334, 1731–1734.

Ko, M., Zou, K., Minagawa, H., Yu, W., Gong, J.-S., Yanagisawa, K., & Michikawa, M. (2005) Cholesterol-mediated neurite outgrowth is differently regulated between cortical and hippocampal neurons. J. Biol. Chem., 280, 42759–42765.

Korinek, M., Vyklicky, V., Borovska, J., Lichnerova, K., Kaniakova, M., Krausova, B., Krusek, J., Balik, A., Smejkalova, T., Horak, M., & Vyklicky, L. (2015) Cholesterol modulates open probability and desensitization of NMDA receptors. J. Physiol., 593, 2279–2293.

Kurzchalia, T.V. & Ward, S. (2003) Why do worms need cholesterol? Nat. Cell Biol., 5, 684–688.

Landrum, M.J., Lee, J.M., Benson, M., Brown, G.R., Chao, C., Chitipiralla, S., Gu, B., Hart, J., Hoffman, D., Jang, W., Karapetyan, K., Katz, K., Liu, C., Maddipatla, Z., Malheiro, A., McDaniel, K., Ovetsky, M., Riley, G., Zhou, G., Holmes, J.B., Kattman, B.L., & Maglott, D.R. (2018) ClinVar: improving access to variant interpretations and supporting evidence. Nucleic Acids Res., 46, D1062–D1067.

Lang, T., Bruns, D., Wenzel, D., Riedel, D., Holroyd, P., Thiele, C., & Jahn, R. (2001) SNAREs are concentrated in cholesterol-dependent clusters that define docking and fusion sites for exocytosis. EMBO J., 20, 2202–2213.

Lee, A.G. (2004) How lipids affect the activities of integral membrane proteins. Biochim. Biophys. Acta, 1666, 62–87.

Li, H. & Papadopoulos, V. (1998) Peripheral-type benzodiazepine receptor function in cholesterol transport. Identification of a putative cholesterol recognition/interaction amino acid sequence and consensus pattern. Endocrinology, 139, 4991–4997.

Linetti, A., Fratangeli, A., Taverna, E., Valnegri, P., Francolini, M., Cappello, V., Matteoli, M., Passafaro, M., & Rosa, P. (2010) Cholesterol reduction impairs exocytosis of synaptic vesicles. J. Cell Sci., 123, 595–605.

Liu, J.-P., Tang, Y., Zhou, S., Toh, B.H., McLean, C., & Li, H. (2010) Cholesterol involvement in the pathogenesis of neurodegenerative diseases. Mol. Cell. Neurosci., 43, 33–42.

Liu, Y., Ross, J.F., Bodine, P.V.N., & Billiard, J. (2007) Homodimerization of Ror2 tyrosine kinase receptor induces 14-3-3(beta) phosphorylation and promotes osteoblast differentiation and bone formation. Mol. Endocrinol., 21, 3050–3061.

Lomize, A.L., Hage, J.M., & Pogozheva, I.D. (2018) Membranome 2.0: database for proteome-wide profiling of bitopic proteins and their dimers. Bioinformatics, 34, 1061–1062.

Lu, W., Yamamoto, V., Ortega, B., & Baltimore, D. (2004) Mammalian Ryk is a Wnt coreceptor required for stimulation of neurite outgrowth. Cell, 119, 97–108.

Maguire, P.A. & Druse, M.J. (1989) The influence of cholesterol on synaptic fluidity, dopamine D1 binding and dopamine-stimulated adenylate cyclase. Brain Res. Bull., 23, 69–74.

Mann, R.K. & Beachy, P.A. (2000) Cholesterol modification of proteins. Biochim. Biophys. Acta, 1529, 188–202.

Martin, M.G., Ahmed, T., Korovaichuk, A., Venero, C., Menchón, S.A., Salas, I., Munck, S., Herreras, O., Balschun, D., & Dotti, C.G. (2014) Constitutive hippocampal cholesterol loss underlies poor cognition in old rodents. EMBO Mol. Med., 6, 902–917.

Martin, M.G., Perga, S., Trovò, L., Rasola, A., Holm, P., Rantamäki, T., Harkany, T., Castrén, E., Chiara, F., & Dotti, C.G. (2008) Cholesterol loss enhances TrkB signaling in hippocampal neurons aging in vitro. Mol. Biol. Cell, 19, 2101–2112.

Martín, M.G., Pfrieger, F., & Dotti, C.G. (2014) Cholesterol in brain disease: sometimes determinant and frequently implicated. EMBO Rep., 15, 1036–1052.

Mauch, D.H., Nägler, K., Schumacher, S., Göritz, C., Müller, E.C., Otto, A., & Pfrieger, F.W. (2001) CNS synaptogenesis promoted by glia-derived cholesterol. Science, 294, 1354–1357.

Maya Vetencourt, J.F., Sale, A., Viegi, A., Baroncelli, L., De Pasquale, R., O’Leary, O.F., Castrén, E., & Maffei, L. (2008) The antidepressant fluoxetine restores plasticity in the adult visual cortex. Science, 320, 385–388.

Michikawa, M. & Yanagisawa, K. (1999) Inhibition of cholesterol production but not of nonsterol isoprenoid products induces neuronal cell death. J. Neurochem., 72, 2278–2285.

Mottaz, A., David, F.P.A., Veuthey, A.-L., & Yip, Y.L. (2010) Easy retrieval of single amino-acid polymorphisms and phenotype information using SwissVar. Bioinformatics, 26, 851–852.

Nieweg, K., Schaller, H., & Pfrieger, F.W. (2009) Marked differences in cholesterol synthesis between neurons and glial cells from postnatal rats. J. Neurochem., 109, 125–134.

Nikoletopoulou, V., Lickert, H., Frade, J.M., Rencurel, C., Giallonardo, P., Zhang, L., Bibel, M., & Barde, Y.-A. (2010) Neurotrophin receptors TrkA and TrkC cause neuronal death whereas TrkB does not. Nature, 467, 59–63.

Niwa, Y.S. & Niwa, R. (2014) Neural control of steroid hormone biosynthesis during development in the fruit fly Drosophila melanogaster. Genes Genet. Syst., 89, 27–34.

Páll, S. & Hess, B. (2013) A flexible algorithm for calculating pair interactions on SIMD architectures. arXiv [physics.comp-ph],.

Park, H. & Poo, M.-M. (2013) Neurotrophin regulation of neural circuit development and function. Nat. Rev. Neurosci., 14, 7–23.

Parrinello, M. & Rahman, A. (1981) Polymorphic transitions in single crystals: A new molecular dynamics method. J. Appl. Phys., 52, 7182.

Pereira, D.B. & Chao, M.V. (2007) The tyrosine kinase Fyn determines the localization of TrkB receptors in lipid rafts. J. Neurosci., 27, 4859–4869.

Peters, M., Katz, B., Lev, S., Zaguri, R., Gutorov, R., & Minke, B. (2017) Depletion of Membrane Cholesterol Suppresses Drosophila Transient Receptor Potential-Like (TRPL) Channel Activity. Curr. Top. Membr., 80, 233–254.

Pfrieger, F.W. (2003a) Role of cholesterol in synapse formation and function. Biochim. Biophys. Acta, 1610, 271–280.

Pfrieger, F.W. (2003b) Outsourcing in the brain: do neurons depend on cholesterol delivery by astrocytes? Bioessays, 25, 72–78.

Pfrieger, F.W. & Ungerer, N. (2011) Cholesterol metabolism in neurons and astrocytes. Prog. Lipid Res., 50, 357–371.

Prasanna, X., Chattopadhyay, A., & Sengupta, D. (2014) Cholesterol modulates the dimer interface of the β2-adrenergic receptor via cholesterol occupancy sites. Biophys. J., 106, 1290–1300.

Prasanna, X., Sengupta, D., & Chattopadhyay, A. (2016) Cholesterol-dependent Conformational Plasticity in GPCR Dimers. Sci. Rep., 6, 31858.

Quan, G., Xie, C., Dietschy, J.M., & Turley, S.D. (2003) Ontogenesis and regulation of cholesterol metabolism in the central nervous system of the mouse. Brain Res. Dev. Brain Res., 146, 87–98.

Rukmini, R., Rawat, S.S., Biswas, S.C., & Chattopadhyay, A. (2001) Cholesterol organization in membranes at low concentrations: effects of curvature stress and membrane thickness. Biophys. J., 81, 2122–2134.

Saito, K., Dubreuil, V., Arai, Y., Wilsch-Bräuninger, M., Schwudke, D., Saher, G., Miyata, T., Breier, G., Thiele, C., Shevchenko, A., Nave, K.-A., & Huttner, W.B. (2009) Ablation of cholesterol biosynthesis in neural stem cells increases their VEGF expression and angiogenesis but causes neuron apoptosis. Proc. Natl. Acad. Sci. U. S. A., 106, 8350–8355.

Suzuki, S., Kiyosue, K., Hazama, S., Ogura, A., Kashihara, M., Hara, T., Koshimizu, H., & Kojima, M. (2007) Brain-Derived Neurotrophic Factor Regulates Cholesterol Metabolism for Synapse Development. Journal of Neuroscience, 27, 6417–6427.

Suzuki, S., Numakawa, T., Shimazu, K., Koshimizu, H., Hara, T., Hatanaka, H., Mei, L., Lu, B., & Kojima, M. (2004) BDNF-induced recruitment of TrkB receptor into neuronal lipid rafts: roles in synaptic modulation. J. Cell Biol., 167, 1205–1215.

Tate, J.G., Bamford, S., Jubb, H.C., Sondka, Z., Beare, D.M., Bindal, N., Boutselakis, H., Cole, C.G., Creatore, C., Dawson, E., Fish, P., Harsha, B., Hathaway, C., Jupe, S.C., Kok, C.Y., Noble, K., Ponting, L., Ramshaw, C.C., Rye, C.E., Speedy, H.E., Stefancsik, R., Thompson, S.L., Wang, S., Ward, S., Campbell, P.J., & Forbes, S.A. (2019) COSMIC: the Catalogue Of Somatic Mutations In Cancer. Nucleic Acids Res., 47, D941–D947.

The UniProt Consortium (2017) UniProt: the universal protein knowledgebase. Nucleic Acids Res., 45, D158–D169.

Waterhouse, A.M., Procter, J.B., Martin, D.M.A., Clamp, M., & Barton, G.J. (2009) Jalview Version 2--a multiple sequence alignment editor and analysis workbench. Bioinformatics, 25, 1189–1191.

Wendler, F., Franch-Marro, X., & Vincent, J.-P. (2006) How does cholesterol affect the way Hedgehog works? Development, 133, 3055–3061.

Zonta, B. & Minichiello, L. (2013) Synaptic membrane rafts: traffic lights for local neurotrophin signaling? Front. Synaptic Neurosci., 5, 9.

